# Trehalose Consumption Ameliorates Pathogenesis in an Inducible Mouse Model of the Fragile X-associated Tremor/Ataxia Syndrome

**DOI:** 10.1101/2023.05.08.539785

**Authors:** Emre Kul, Oliver Stork

**Affiliations:** Department of Genetics & Molecular Neurobiology, Institute of Biology, Otto-von-Guericke University Magdeburg, Germany; Center for Behavioral Brain Sciences, Magdeburg, Germany

**Keywords:** FXTAS, Repeat Expansion, Neurodegeneration, Cerebellum, Motor Behavior, Trehalose, Autophagy, Transgenic Mouse

## Abstract

Trehalose is a naturally occurring sugar found in various food and pharmaceutical preparations with the ability to enhance cellular proteostasis and reduce formation of toxic intracellular protein aggregates, making it a promising therapeutic candidate for various neurodegenerative disorders. Here, we explored the effectiveness of nutritional trehalose supplementation in ameliorating symptoms in a mouse model of Fragile X-associated tremor/ataxia syndrome (FXTAS), an incurable late onset manifestation of moderately expanded trinucleotide CGG repeat expansion mutations in the 5’ untranslated region of the fragile X messenger ribonucleoprotein 1 gene (*fmr1*). An inducible mouse model of FXTAS expressing 90 CGG repeats in the brain had been previously developed, which faithfully captures hallmarks of the disorder, the formation of intracellular inclusions and the disturbance of motor function. Taking advantage of the inducible nature of the model, we investigated the therapeutic potential of orally administered trehalose under two regimens, modelling disease prevention and disease treatment. Trehalose’s effectiveness in combating protein aggregation is frequently attributed to its ability to induce autophagy. Accordingly, trehalose supplementation under the prevention regimen ameliorated the formation of intranuclear inclusions and improved the motor deficiencies resulting from the induced expression of 90 CGG repeats, but it failed to reverse the existing nuclear pathology as a treatment strategy. Given the favorable safety profile of trehalose, it is promising to further explore the potential of this agent for early stage FXTAS.

## INTRODUCTION

Trehalose (PubChem CID: 7427) is a naturally occurring non-reducing disaccharide sugar composed of two glucose molecules. It is produced by bacteria, plants, yeast, fungi, and invertebrates but not by mammals. Trehalose is present in various food sources including honey, mushrooms, and shellfish, and used as a glucose or sucrose substitute in various food products (Richards et al. 2002; Moriano and Alamprese 2017). Trehalose is also known as a cryoprotectant to the freeze-dried food industry and as a stabilizer excipient for pharmaceuticals due to its ability to preserve proteins and lipids under various stress conditions (Lee et al. 2013; Stefanello et al. 2018). Importantly, the favorable use of trehalose extends to therapeutics research. Trehalose was shown to have beneficial effects in clearing mutant proteins, decreasing aggregates, improving motor function, and extending lifespan in preclinical studies of repeat expansion disorders such as Huntington’s disease (Tanaka et al. 2004), amyotrophic lateral sclerosis (ALS) (Li et al. 2015), and spinocerebellar ataxia type 3 (SCA3) (Santana et al. 2020). Owing to its preclinical therapeutic potential, trehalose has now been brought to the clinic under the designation SLS-005, as an investigational drug to treat ALS (clinical trial database identifier: NCT05136885), SCA3 (NCT05490563), Alzheimer’s Disease (NCT05332678), and Parkinson’s Disease (NCT05355064).

Furthermore, trehalose has received orphan drug designation from the United States Food and Drug Administration (FDA) for the treatment of ALS (accessdata.fda.gov/scripts/opdlisting/oopd/detailedIndex.cfm?cfgridkey=781920) and SCA3 (accessdata.fda.gov/scripts/opdlisting/oopd/detailedIndex.cfm?cfgridkey=442014).

Fragile X-associated tremor/ataxia syndrome (FXTAS) shares significant similarities in pathology and phenotype with other neurodegenerative disorders that are currently under investigation for the therapeutic potential of trehalose. Despite substantial efforts to develop therapeutic strategies, including screening FDA-approved compounds as well as targeted approaches, FXTAS currently lacks any effective treatments. Patients only receive supportive care, aimed at managing their individual symptoms that cause the most significant problems (Sahdeo et al. 2014; Hagerman and Hagerman 2016; Reddy et al. 2020). FXTAS is the neurodegenerative member of a family of X chromosome-linked disorders that stem from trinucleotide CGG repeat expansions (CGG^exp^) in the 5’ untranslated region (UTR) of the fragile X messenger ribonucleoprotein 1 (*fmr1*) gene. The disorder affects 1 in 4000 men above the age of 55 and is caused by fmr1 premutation alleles that harbor expansions of 55-200 CGG repeats. While 40% of male carriers develop FXTAS, random X chromosome inactivation greatly reduces the prevalence in females. Furthermore, individuals carrying premutation alleles have an increased risk of transmitting a full fmr1 mutation to their offspring, leading to the development of Fragile X Syndrome in subsequent generations (Hagerman and Hagerman 2016; Wheeler et al. 2017).

FXTAS is characterized by tremor and cerebellar ataxia with a neuropathology including white matter lesions and generalized atrophy. Ubiquitin-positive intranuclear inclusions found in neurons and glia are a hallmark of the disorder and is believed to represent a toxic mechanism of action. Although the molecular basis of FXTAS is not completely understood, three potentially druggable pathomechanisms have been proposed. A well-evidenced mechanism involves RNA toxic gain-of-function by sequestration of RNA-binding proteins (RBPs) via interactions with the enlarged hairpin structure that fmr1 premutation mRNA harbors. This can drive formation of intranuclear inclusions due to aggregation of the RBPs and their downstream targets. The functional depletion of the aggregating RBPs is believed to interfere with cellular homeostasis and cause toxicity. A second proposed mechanism involves non-canonical translation of the 5’ UTR of the premutation mRNA into homopolypeptides via repeat-associated non-AUG (RAN) translation. RAN translation can initiate from a near-cognate codon upstream of CGG^exp^ and give rise to homopolyglycine known as FMRpolyG that is prone to aggregation. The translated FMRpolyG can initially form small cytoplasmic aggregates and can be detected in FXTAS intranuclear inclusions. The third and relatively less investigated molecular mechanism involves the formation of R-loop structures containing RNA:DNA hybrids formed between the CGG^exp^ mRNA transcript and its DNA template. R-loops may cause local DNA damage and cellular toxicity and induce repeat-length changes during DNA replication leading to genome instability (Hagerman and Hagerman 2016; Zhao and Usdin 2016; Gohel et al. 2019).

In this study, we explored the therapeutic potential of orally administered trehalose in mitigating FXTAS-elated pathology and behavioral impairments in a well-established mouse model expressing a trinucleotide repeat tract of 90 CGGs in the brain upon induction with doxycycline (dox) (hereafter referred to as P90CGG). This is a powerful mouse model of FXTAS that recapitulates the inclusion pathology as well as the motor phenotype of the disorder (Hukema et al. 2015; Castro et al. 2017). To explore the efficacy of trehalose in preventing disease onset or treating established phenotypes, we administered trehalose to the mice through their drinking water during or after CGG^exp^ transgene induction, respectively. As a preventive strategy, trehalose limited the aggregation and ameliorated the motor deficits.

## METHODS

### Animals

Inducible transgenic animals (referred to as PrP-rtTA/TRE-90xCGG or P90CGG for short) were obtained by crossing single transgenic mutants carrying a 90CGGrepeat tract along with a tetracycline responsive element (TRE-90CGG) with the transgenic prion protein-reverse tetracycline transactivator (PrP-rtTA) driver line, as described previously (Hukema et al. 2015). Upon induction with dox, these mice express a 90 CGG trinucleotide tract in the brain as dictated by the PrP promoter. Wild type C57BL/6JBomTac (BL6) mice were purchased from M&B Taconic. All mice were bred and maintained at the Institute of Biology, Otto-von-Guericke University Magdeburg, Germany under standard laboratory conditions with inverted 12-hour dark/light cycle. Mice were weaned four weeks after birth, group-housed, and provided with *ad libitum* food (R/M-H V-1534 or R/M-H-A153-D04004+4600 mg/kg doxycycline hyclate (Ssniff)) and water (plain or containing 2% v/v food grade trehalose (FormMed)). Tail biopsies were collected for genotyping and earmarks were made for mouse identification. Genotyping was performed as previously described (Hukema et al. 2015). All experimental procedures were approved by local ethics committee Landesverwaltungsamt Sachsen-Anhalt (CEEA# 42502-2-1219UniMD) and met the guidelines of local and European regulations (European Union directive no. 2010/63/EU). Animal experiments were performed during the active phase of the mice.

### Trehalose Administration

#### Preventive Strategy

To fight RAN translation driven aggregation as it occurs, food grade trehalose was orally supplemented *ad libitum* to dox-inducible P90CGG mice for 12 weeks after weaning, with trehalose and dox (*ad libitum* in feed) supplementation simultaneously starting and ending. Following behavioral testing at age of ∼16 weeks, brain tissue was collected.

#### Treatment Strategy

To reverse already developed phenotypes, oral trehalose was provided *ad libitum* for 12 weeks to dox-inducible P90CGG mice after the 12-week dox induction period had ended, and brain tissue was collected after behavioral testing at ∼28 weeks of age.

### Rotarod

The motor performance of mice was assessed using the Rotarod test (Ugo Basile), over 3 days (Derbis et al. 2021). Mice were trained on the rotating rod at 15 rpm constant speed for a maximum of 60 s in four trials, and then tested on the rod rotating at various constant speeds (15-40 rpm) in two trials for each speed, recording the latency to fall off the rod. On the final day, mice were tested in four 5 min trials with accelerating rotation from 4 to 40 rpm.

### Tissue Processing and Immunofluorescence

Mice were anesthetized with isoflurane and sacrificed by decapitation, and the brain tissue was immediately removed. The right hemispheres were drop fixed in 4% PFA/PBS at 4 °C overnight, then cryoprotected in 30% sucrose/PBS containing 0.02% sodium azide, and embedded in O.C.T. compound (Sakura) before being frozen in liquid nitrogen cooled isopentane (Sigma). The frozen blocks were cryostat-sectioned into 7 μm sagittal sections (Leica). Sections were mounted on poly-L-lysine coated glass slides, air-dried, and stored at 4°C. Sections were then subjected to heat-induced epitope retrieval in 0.01M sodium citrate (pH=6.0) and proteolytic-induced epitope retrieval with 5 μg/ml proteinase K (Roth) at 37°C. After blocking for endogenous biotin and mouse Ig (Vector), the sections were incubated with 8FM anti-FMRpolyG antibody (1:200) (kind gift from Nicholas Charlet-Berguerand) and biotinylated secondary antibody (Vector), followed by visualization with Cyanine 5-conjugated streptavidin (Invitrogen). Finally, sections were counterstained with DAPI and covered with Immu-Mount (Shandon).

### FMRpolyG Foci Analysis

Cerebellum lobule X photomicrographs were taken with an epifluorescence microscope (Leica) at 630x magnification, with a z-axis step size of 2 μm. Open-source image processing software Fiji was used to analyze the images. A custom cell counter script was developed to randomly display cropped images of 30 × 30 μm generated from whole photomicrographs. 400 nuclei stained with DAPI in the granular layer of lobule X were counted per mouse, and for each nucleus, a researcher blind to the treatment groups recorded whether it contained an FMRpolyG+ intranuclear inclusion, and measured the inclusion’s longitudinal size, as described previously (Derbis et al. 2021). The data were used to calculate a percentage value and an average size value for each mouse.

### Western Blot

Cerebellum tissue was lysed on ice with a Dounce homogenizer (Wheaton) in lauryl maltoside lysis buffer (1% n-dodecyl-beta-D-maltoside, 1% NP-40, 1 mM Na_3_VO_4_, 2mM EDTA, 50 mM Tris-HCl, 150 mM NaCl, 0.5% deoxycholic acid, 1 mM AEBSF, 1 μM pepstatin A, 1 mM NaF, 1 tablet Pierce protease inhibitor (Thermo Fisher)). Homogenized samples were agitated on a rotator at 40 rpm at 4°C for 30 min. Samples were then centrifuged for 30 min at 4°C at 16000 rcf. Protein concentration of each sample was determined using the Bradford assay (Bio-Rad). Protein samples were separated by SDS-PAGE (15%) and transferred to PVDF membranes (Immobilon-FL, Millipore). Membranes were incubated with Intercept Blocking Buffer (Li-COR), then with primary antibodies (anti-LC3 A/B-12741, Cell Signaling, 1:1500 and anti-alpha tubulin-T6199, Sigma, 1:10000), followed by fluorescent secondary antibody incubation. Blotted membranes were scanned with the Odyssey scanner (LI-COR) and analyzed using Odyssey ImageStudio software (LI-COR).

### Statistical Analysis

All statistical analyses were conducted in Prism software (GraphPad). First, data were checked for normality using Shapiro-Wilk normality test. For normally distributed data, Student’s t-test or one-way ANOVA was performed, while non-normally distributed data were analyzed using Mann-Whitney U-test or Kruskal-Wallis test. Two-way ANOVA was used for comparing two factors, and repeated measures two-way ANOVA was used when repeat measures were available for one of the factors. For analyses involving three factors, including cases where one factor is a repeated measures factor, data were analyzed using three-way ANOVA. Statistically significant treatment effects and factor interactions resulting from hypothesis testing were reported for ANOVA analyses. Post-hoc comparisons were conducted with Fisher’s LSD test when applicable. A p-value of less than 0.05 was considered statistically significant for all analyses.

## RESULTS

To investigate the therapeutic potential of trehalose in counteracting the pathological processes as they occur, we administered 2% v/v oral trehalose (treh) to P90CGG mice during the CGG^exp^ induction period and we refer to this as preventive timeline. In the treatment timeline, we investigated the possibility of reversal of already inflicted damage by the administration of trehalose following the end of CGG^exp^ induction period.

### Preventive trehalose limits aggregation and ameliorates motor behavior

Intranuclear inclusions are found throughout the brain of the P90CGG mice after an induction period of 12 weeks. In the cerebellum, they are most numerous in granule cell layer of the lobule X (Fig.1B) and stain positively using an antibody raised against FMRpolyG homopolypeptide (Fig.1C) (Derbis et al. 2021). Therefore, upon 12-week of dox induction (Fig.1A), we quantified the number and size of these inclusions from the lobule X of treh-supplemented and non-supplemented mice (N(dox+.treh-)=11, N(dox+.treh+)=11). The proportion of nuclei with detectable FMRpolyG foci was significantly decreased in the treh-supplemented group (Fig.1D; Student’s t-test, t=3.154, df=20, P= 0.0050). The average size of the FMRpolyG foci was also reduced upon treh supplementation (Fig.1E; Student’s t-test, t=2.832, df=20, P= 0.0103).

**Figure 1.**
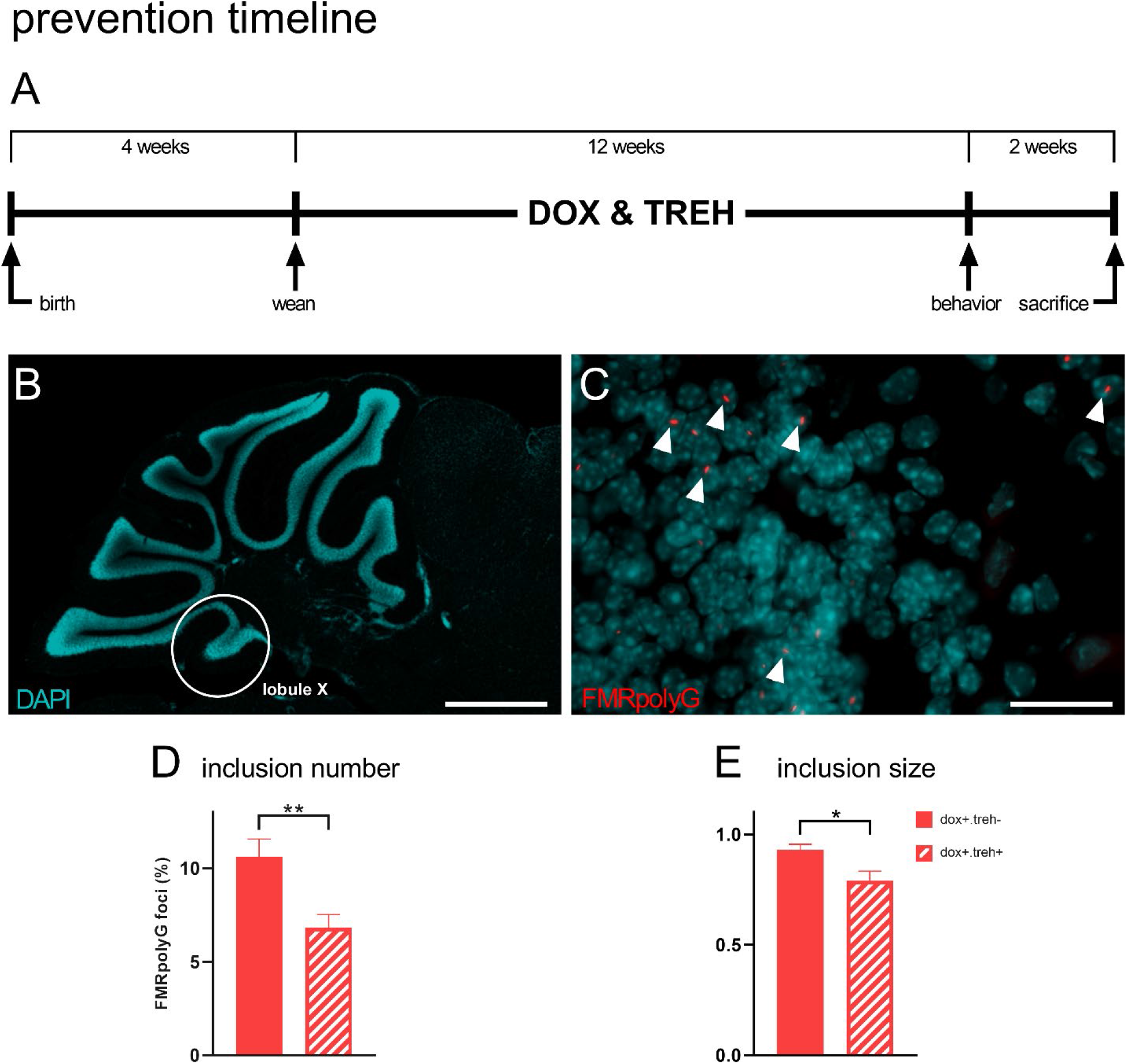
Trehalose prevention timeline results in decreased inclusion load in cerebellum lobule X. **A**. Timeline showcasing the dox-induction and trehalose supplementation schedules used for the prevention strategy. **B**. Representative photomicrograph of a sagittal section of mouse cerebellum indicating the position of lobule X (white contour), the readout region for inclusion quantifications. Scale bar: 1 mm. **C**. High magnification photomicrograph of granule cell layer of the lobule X showing the immunostained FMRpolyG-positive inclusions (red) colocalizing with various nuclei (blue: DAPI) within the granule cell layer. White arrowheads: FMRpolyG foci (not all foci marked). Scale bar: 25 μm. **D**. Trehalose supplementation had significantly reduced the number nuclei with FMRpolyG foci in the dox-induced P90CGG mice. **E**. The FMRpolyG foci were significantly smaller in size with the trehalose-supplemented group in comparison to treh-group. Data presented as mean ±S.E.M. *p<0.05, **p<0.01.

We evaluated four groups of mice (N(dox-.treh-)=10, N(dox-.treh+)=10, N(dox+.treh-)=11, N(dox+.treh+)=11) for motor performance and potential deficits on the Rotarod under our preventive timeline. During the training session performed at low speed (15 rpm), no significant effects of either dox or treh were observed (Fig.2A; 3-way ANOVA, dox-effect; F(1,38)=2.506, n.s., treh-effect; F(1,38)=2.098, n.s.) and all four groups reached a comparable acclimation level at the end of the session. However, performance deficits became evident when mice were tested with increased rotation speed, as the impairing effects of transgene induction were present at 31rpm and higher, and were efficiently counteracted by treh-supplementation (Fig.2B; 3-way ANOVA, dox-effect; F(1,38)=26.39, P<0.0001, treh-effect; F(1,38)=6.482, P=0.0151, dox x treh interaction; F(1,38)=6.521, P=0.0148). When the same animals were finally tested on a steadily accelerating rod from 4 to 40 rpm in 5 minutes, the same negative effect of transgene induction on the latencies to fall off the rod and recovery of performance by treh-supplementation was observed (2-way ANOVA, dox-effect; F(1,38)=24.77, P<0.0001, treh-effect; F(1,38)=10.66, P=0.0023, dox x treh interaction; F(1,38)=7.622, P=0.0088). Moreover, a significant negative correlation between the abundance of inclusions in lobule X and motor performance on the accelerating Rotarod was observed (Fig.2D; Pearson correlation, r=-0.4457, P=0.0376).

**Figure 2.**
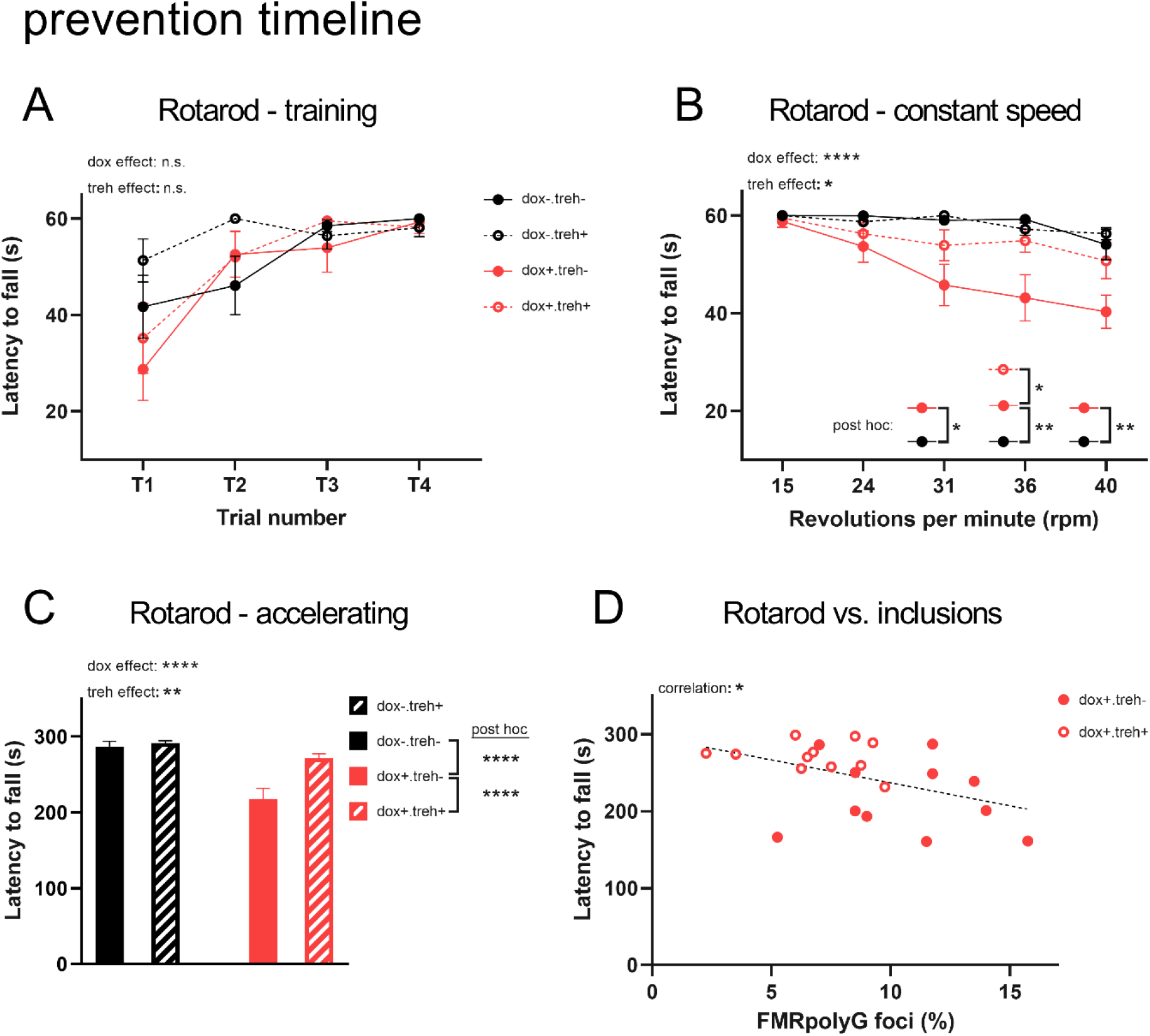
Preventive trehalose recovers the deficits in motor performance resulted from dox-induction. **A**. No significant effects were observed either for dox or treh in the latency to fall during the 4-trial 15-rpm training session of the Rotarod test. **B**. During the constant speed test, dox-induction significantly decreased the latencies of P90CGG mice to fall off the rod whereas trehalose significantly increased the time the mice spent on the rod rotating at various speeds. **C**. Dox-induction significantly decreased the latency to fall when the mice were tested in a speed ramp setting during the accelerating rotarod test. Contrastingly, trehalose significantly increased the latency of the mice to fall off the rod. **D**. A statistically significant negative correlation was present between the latencies to fall off the rod of individual mice during the accelerating rotarod task and the number of nuclei with FMRpolyG foci in the granule cell layer of the lobule X. Data presented as mean ±S.E.M. *p<0.05, **p<0.01, ****p<0.0001, ^n.s.^p>0.05.

No significant effects of either dox of treh has been found on the body weights of mice (Fig.S2A; 2-way ANOVA, dox-effect; F(1,38)=2.352, n.s., treh-effect; F(1,38)=0.9545, n.s.). Moreover, to control for genotype-independent effects dox and treh may potentially have on motor phenotype, wild-type black-6 (BL6) mice were subjected to 12-week dox or treh supplementation (Fig.S1A) and tested on Rotarod (N(water_BL6_)=8, N(treh_BL6_)=9, N(dox_BL6_)=8). No effect of these treatments were observed during the training session (Fig.S1B; 2-way RM ANOVA, substance-effect; F(2,22)=1.138, n.s.), constant speed session (Fig.S1C; 2-way RM ANOVA, substance-effect; F(2,22)=1.129, n.s.) or acceleration session (Fig.S1D; Kruskal-Wallis test, K-W=1.711, n.s.). Again, body weights did not differ between control, dox, and treh groups (Fig.S1E; 1-way ANOVA, substance-effect; F(2,22)=2.535, n.s.).

### Trehalose increased the levels of the autophagy marker LC3-II in the P90CGG cerebella

Trehalose has several previously described properties that can lead to a removal of aggregates and neuroprotection. Among these, the best investigated mechanism is through the induction of autophagy (Mardones et al. 2016). To identify whether the positive effects we observed may be linked to autophagy, we performed western blot analysis and quantified the levels of an autophagy marker, LC3-II, from cerebellum tissue whole protein extracts. LC3-II, an autophagosome associated protein, which results from posttranslational modification of LC3 was evaluated as a measure of autophagy induction (Tanida and Waguri 2010). Four experimentally naïve groups of mice (N(dox-.treh-)=8, N(dox-.treh+)=8, N(dox+.treh-)=8, N(dox+.treh+)=8) under the preventive timeline were used with tissue collection done during continued treh supplementation. Independent of dox induction, treh increased the levels of LC3-II in the mouse cerebellum (Fig.3A-B; 2-way ANOVA, treh-effect; F(1,28)=5.655, P=0.0245, dox-effect; F (1,28)=0.02427, n.s., dox x treh interaction; F (1,28)=0.01815, n.s.).

**Figure 3.**
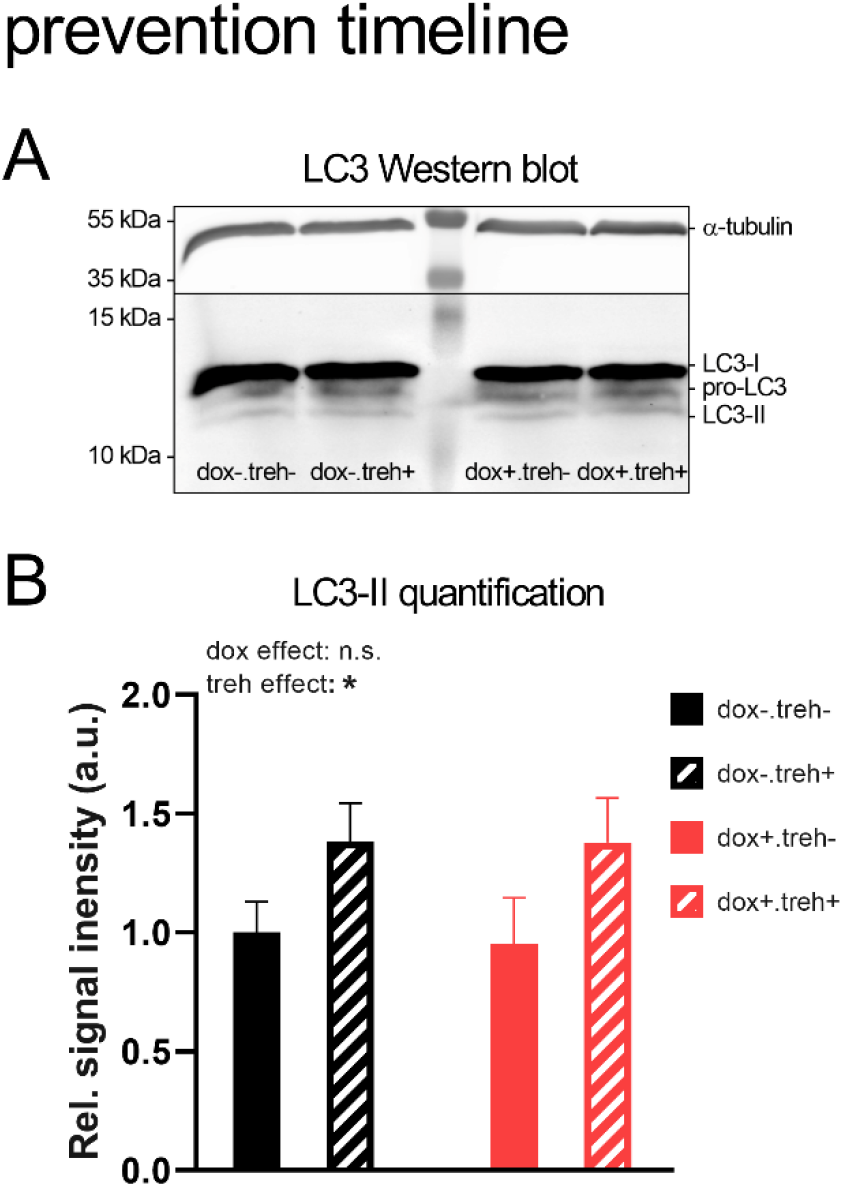
Trehalose increased the LC3-II levels in the cerebellum of P90CGG mice. **A**. Representative western blot scan showing fluorescent signal specific to α-tubulin (upper segment) and LC3 (lower segment) under four different dox and trehalose conditions. Notice the three isoforms of LC3 detected by the primary antibody. **B**. Quantification of the LC3-II signal (lowest molecular weight band) normalized to α-tubulin signal. Trehalose significantly increased the LC3-II signal, whereas dox-induction had no effect. Data presented as mean ±S.E.M. *p<0.05, ^n.s.^p>0.05.

### Trehalose treatment fails to revert pathology and motor deficits

To investigate the potential of trehalose for reverting already developed FXTAS phenotypes, four groups of P90CGG mice (N(dox-.treh-)=10, N(dox-.treh+)=9, N(dox+.treh-)=10, N(dox+.treh+)=9) were subjected to the treatment timeline (Fig.4A). In the lobule X tissue of dox-induced groups, we furthermore quantified inclusion number and inclusion size (N(dox-.treh+)=9, N(dox+.treh+)=8, with one sample excluded due to tissue damage during processing). We observed no differences in the number (Fig4B; Student’s t-test, t=0.09053, df=16, n.s.) or in the size (Fig4C; Student’s t-test, t=0.04328, df=16, n.s.) of inclusions. Further, as before, no effects of either dox-induction or treh-supplementation were observed during the training session with all groups reaching similar acclimation levels (Fig.4D; 3-way ANOVA, dox-effect; F(1,34)=2.649, n.s., treh-effect; F(1,34)=1.472, n.s.). During the constant speed Rotarod test, transgene-induction impaired motor performance as expected, but no effect of treh could be established (Fig.4E; 3-way ANOVA, dox-effect; F(1,34)=15.56, P=0.0004, treh-effect; F(1,34)=0.8237, n.s). The detrimental effect of transgene-induction persisted in the accelerating ramp test with trehalose again failing to recover performance (Fig.4F; 2-way ANOVA, dox-effect; F(1,34)=10.99, P=0.0022, treh-effect; F(1,34)=1.011, n.s.). Dox-induced mice were significantly heavier than their counterparts; again trehalose had no effect (Fig.S2B; 2-way ANOVA, dox-effect; F(1,34)=35.81, P<0.0001, treh-effect; F(1,34)=1.278, n.s.). To control for potential bias in motor performance due to the change in body weights, we have performed a correlation analysis of body weights of dox-induced mice against their acceleration test performance values, but no correlation could be identified (Fig.S2C); Pearson correlation, r=-0.2873, n.s.).

**Figure 4.**
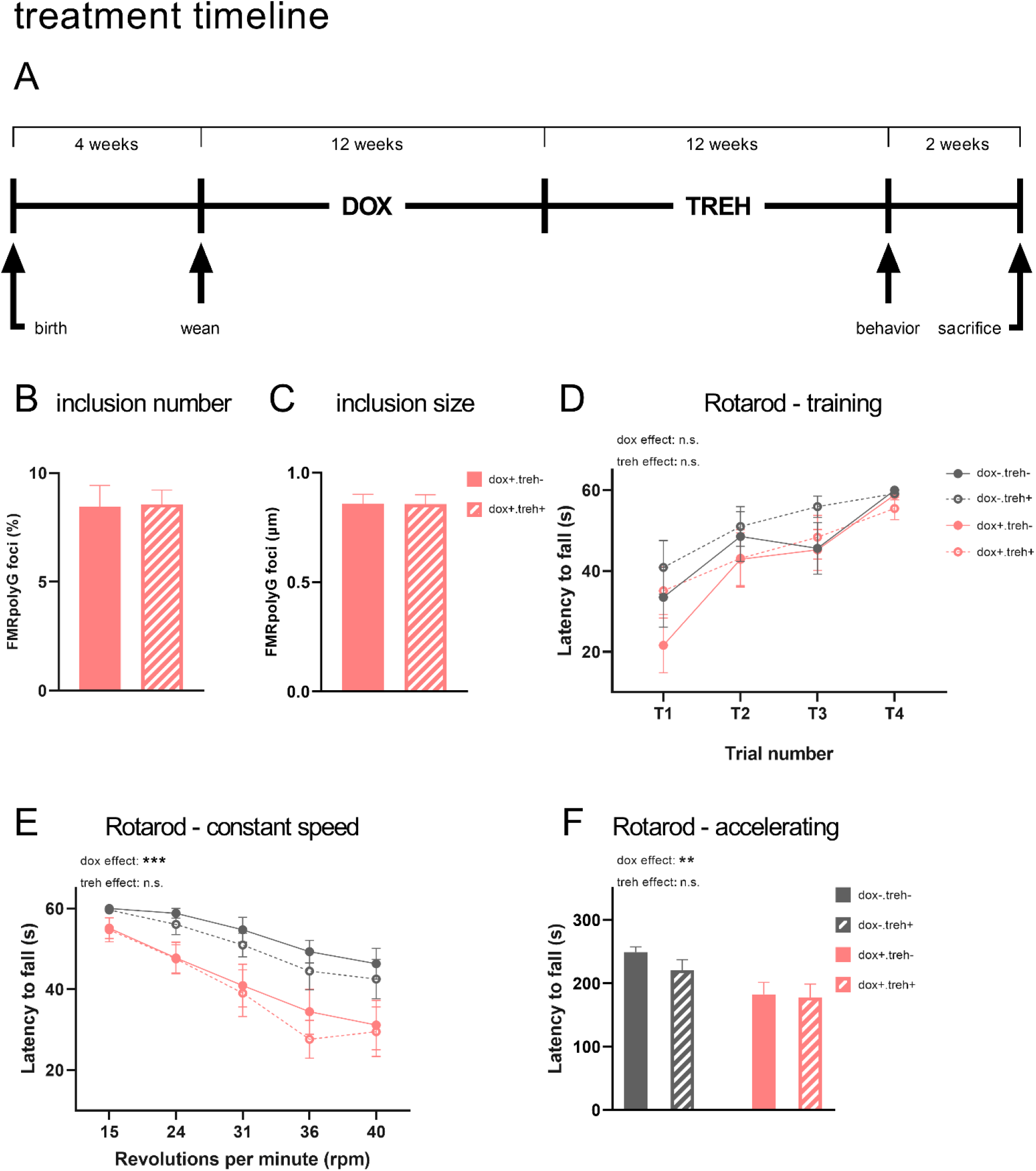
Trehalose is ineffective against pathology and motor deficits as a treatment strategy. **A**. Timeline showcasing the dox-induction and trehalose supplementation schedules used for the treatment strategy. **B**. Trehalose supplementation did not alter the number nuclei with FMRpolyG foci in the dox-induced mice. **C**. The FMRpolyG foci size remained unchanged between the trehalose-supplemented group and treh-group. **D**. No significant effects were observed either for dox or treh in the latencies to fall off the rod during the training session of the Rotarod test. **E**. Dox-induction has significantly reduced the time P90CGG mice remained on the Rotarod during the constant speed test. No statistically significant effect was present for trehalose. **F**. Dox has decreased the latencies to fall off the rod, whereas trehalose had no effect during the acceleration test. Data presented as mean ±S.E.M. **p<0.01, ***p<0.001, ^n.s.^p>0.05.

## DISCUSSION

In this study, we investigated the therapeutic potential of trehalose in preventing and reversing pathological features in the inducible PrP.90xCGG (P90CGG) mouse model of FXTAS. This model recapitulates many of the aspects of the disorder including intranuclear inclusion pathology and motor dysfunction and allows to control the time point of pathology onset (Hukema et al. 2015). It provides a framework to draw parallels to the human condition and has been previously taken advantage of as a testing platform for therapeutic interventions (Derbis et al. 2021). We have previously shown that 12 weeks of dox induction results in readily detectable inclusion pathology in the cerebella of these mice and motor performance deficits on the Rotarod (Castro et al. 2017). A prevention strategy, in which trehalose was supplemented in coincidence with dox induction led to significantly reduced numbers of intranuclear inclusions in the granule cell layer of the cerebellum lobule X and recovered motor performance on Rotarod. By contrast, a treatment timeline of trehalose administration succeeding dox induction was ineffective against either phenotype.

Trehalose is a known autophagy inducer and its beneficial effects in neurodegenerative protein aggregation models have been attributed to this property. Autophagy is typically triggered by the lack of nutrients under the inhibitory control of a nutrient sensor, mechanistic target of rapamycin (mTOR), promoting a degradative recycling action to increase nutrient availability. The same action also favors ridding the cell of misfolded proteins and aggregations. Trehalose, as an mTOR-independent activator, achieves a similar effect by blocking glucose transport, creating an adenosine triphosphate (ATP) deprived starvation-like state within the cell (Mardones et al. 2016). Upon oral intake, trehalose can be detected in brain (Howson et al. 2019) and it has several other properties other than being an autophagy inducer that may provide beneficial effects in its applications concerning neurodegeneration. Trehalose is believed to act against aggregation through its chemical chaperone properties, stabilizing protein structures and their folding state. Trehalose is also known for its antioxidant and anti-inflammatory properties and it may protect cells from degeneration by stimulating enzymatic antioxidants and scavenging reactive oxygen species (Gao et al. 2018; Mizunoe et al. 2018). An indirect gut-brain-axis function has also been hypothesized for trehalose supported by its property of remodeling gut microbiota (Buckley et al. 2021) and its ability to induce autophagy in the brain upon oral intake but not after intraperitoneal injection (Tanji et al. 2015; Lee et al. 2018).

Although dysfunctions in the ubiquitin proteasome system were described for FXTAS, to which the aggregation phenotype was attributed to, more recent reports support a role of maladaptive and dysfunctional autophagy. In fact, FXTAS intranuclear inclusions contain ubiquitin and p62/SQSTM1, two adapter proteins autophagy takes advantage of to target a broad range of cargo from proteins to damaged organelles for degradation (Ma et al. 2019). p62/SQSTM1 and ubiquitin-tagged cargo are associated with LC3 proteins that are autophagosome coat proteins involved in its formation. Accumulation of p62 correlates with stalled autophagic processes (Klionsky et al. 2016), indicating an inability to cope with the increased aggregation. Autophagosome encapsulated cargo can be degraded in bulk, thus aggregated proteins are good candidates for autophagic targeting. Indeed, even large cytoplasmic inclusion bodies can be degraded via autophagy (Pankiv et al. 2007). Due to their important role in the autophagic processes, LC3-II and p62 are also commonly used autophagy markers. Since the levels of LC3-II correlate with formation of mature autophagosomes and reflect autophagy progression (Klionsky et al. 2016), increased levels in the cerebella of the trehalose supplemented P90CGG mice indicate a trehalose-mediated autophagic activation (Fig.3). Importantly, rescue of pathologies through trehalose-mediated autophagic induction has been shown even in systems with initially overwhelmed and dysfunctional autophagy (Sergin et al. 2017).

The decrease in intranuclear inclusion pathology in the trehalose supplemented P90CGG mice during the prevention timeline is consistent with autophagic degradation of FMRpolyG kicking in as it starts to aggregate in the cytoplasm. On the other hand, trehalose was ineffective in reversing the already present pathology and associated motor behavior deficits even in the absence of sustained CGG^exp^ expression. This, in turn, is also not completely unexpected if the drug exerts its effect through autophagy. Since autophagy is a cytoplasmic process, after inclusions are fully formed in the nucleus, they become inaccessible to autophagic degradation especially in post-mitotic cells, where nuclear membrane presents a permanent barrier. Thus, autophagic targeting may also explain absence of FXTAS inclusions in rapidly dividing cells like fibroblasts (Garcia-Arocena et al. 2010). However, these do not exclude additive effects that may play a role due to other properties of trehalose, such as protein structure stabilization.

To make our deductions with respect to effectiveness of trehalose on inclusion pathology, we quantified intranuclear inclusions from the granule cell layer of the cerebellum lobule X. This region is the most impacted tissue by the inclusion pathology in the P90CGG model and was similarly used in our previous study to evaluate effectiveness of therapeutic strategies (Derbis et al. 2021). We found that the motor performance on the Rotarod is inversely correlated with the number of inclusions in this region, suggesting that the improved behavioral outcome upon trehalose supplementation may reflect an amelioration of toxicity and aggregation pathology. It should be considered that other cerebellar regions than lobule X are also involved in Rotarod performance and that our quantifications are based on the condition of the most adversely affected structure of the cerebellum. Thus, the overall functional state of the cerebellum may be better than what we are detecting at the inclusion pathology level from lobule X alone. This may explain why the more remarkable improvement in the motor performance is supported by a moderate (∼35%) decrease in the number of inclusions in lobule X.

From a translational perspective, it has to be considered that P90CGG is an artificial model overexpressing CGG^exp^ outside of the context of *fmr1* (Hukema et al. 2015) and the differential pattern of inclusion frequency between various tissues of the cerebellum may not faithfully recapitulate the human case. The model shows robust pathology, but potential interactions between FXTAS genetic makeup and normal ageing of the organism are not captured in our study. Furthermore, the overexpression of CGG^exp^ in our model may be contributing to the incomplete rescue of the inclusion pathology. Whether a stronger effect of trehalose might be found under slower aggregation conditions, i.e. lower transgene expression, would be interesting to explore. On the other hand, any strategy that targets the RAN translation product, FMRpolyG alone, will be unable to counteract the CGG^exp^ mRNA-mediated toxicity in the nucleus, such as protein sequestration or R-loop formation. Currently, there is no consensus on which molecular mechanism plays the dominant role in driving the FXTAS phenotypes and potential therapeutic targets remain to be largely different for each mechanism (Hagerman and Hagerman 2016). Therefore, it is not possible to exclude the potential contribution of other proposed pathomechanisms to the remaining inclusion pathology following our trehalose administration strategy. Nevertheless, the profound rescue by trehalose surrounding pathology and behavior are promising findings warranting further investigation.

Based on accumulated evidence from preclinical studies associated with repeat expansion and other neurodegenerative disorders, trehalose is currently being evaluated at clinical trials for the treatment of amyotrophic lateral sclerosis, spinocerebellar ataxia type 3, Alzheimer’s disease, and Parkinson’s disease. The pathomechanisms of FXTAS share significant similarities with that of these disorders and the high safety profile of trehalose may hold potential for clinical trials in FXTAS as well. Although the results of our current preclinical study show that a reversal of the already inflicted damage was not possible, the positive effects seen under the preventive timeline are noteworthy, particularly given the late onset nature of FXTAS. Moreover, it will be interesting to combine trehalose with treatments inhibiting nuclear import of FMRpolyG or stimulating its nuclear export that could make aggregates forming in the nucleus accessible for autophagy.

## ACKNOWLEDGEMENTS

We are grateful to F. Blitz, A. Koffi von Hoff, and S. Stork for expert technical assistance and A. Bohnstedt and D. Al-Chakmakchi for devoted and impeccable animal care.

This work was supported by the European Union under grant 01GM1505 ERARE; DrugFXSPreMut and by the German Research Foundation under 3038041027; STO488/9-1 (RU5228 Syntophagy) and 362321501 (RTG 2413 SynAGE) to O.S.

## DATA AVAILABILITY

The data that support the findings of this study are available from the corresponding author, O.S., upon reasonable request.

## BIOGRAPHICAL NOTE

Emre Kul is a neurobiologist with a background in chemistry. He obtained his Ph.D. degree from the Otto-von-Guericke-University Magdeburg, Germany, where he conducted research on mouse models of cerebellar neurodegeneration. His work focused on developing therapeutic strategies for Fragile X-associated tremor/ataxia syndrome (FXTAS), including the use of antisense oligonucleotide therapy and other FDA-approved compounds. Currently, Emre is dedicated to developing novel, high-capacity gene delivery vectors and investigates their applications in gene therapy approaches for monogenic cerebellar ataxias.

Oliver Stork is a neurobiologist with a background in behavioural and molecular neurobiology. He obtained his Ph.D. from the Swiss Federal Technical University Zürich in 1997 and is Professor for Molecular Neurobiology at the Otto-von-Guericke-University Magdeburg since 2008. His research is devoted to the understanding of molecular mechanisms that underlie information storage in the central nervous system and the control of behaviour. In particular, he focusses on the processes in brain circuits that mediate the translation of specific neuronal activity patterns into lasting morphological and neurochemical changes and their relevance for learning and memory, stress, and ageing.

