## Supplemental Figures for "Trehalose Consumption Ameliorates Pathogenesis in an Inducible Mouse Model of the Fragile X-associated Tremor/Ataxia Syndrome"

SUPPLEMENTARY FIGURES

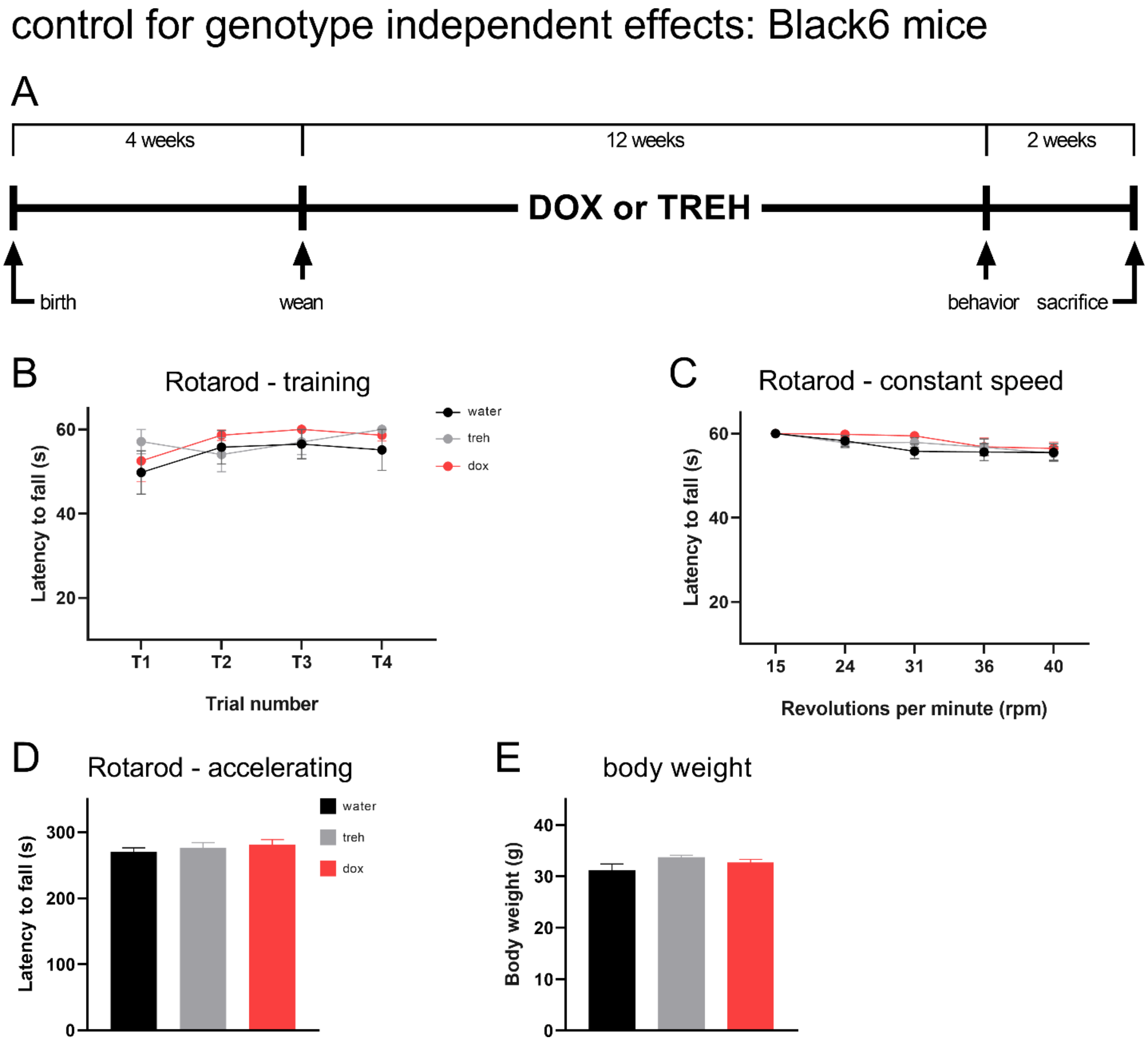


**Supplementary Figure 1. Neither dox nor trehalose showed genotype independent effects altering motor performance or body weight**

**A.** Timeline showcasing the dox-induction and trehalose supplementation schedules used with wild-type BL6 mice to investigate genotype-independent effects of dox and trehalose on motor behavior. BL6 mice supplemented with either dox or trehalose did not differ from plain water drinking counterparts in any of the Rotarod sessions; **B.** training, **C.** constant speed or **D.** acceleration. **E.** No significant changes in the body weights detected between the three groups. Data presented as mean ±S.E.M.

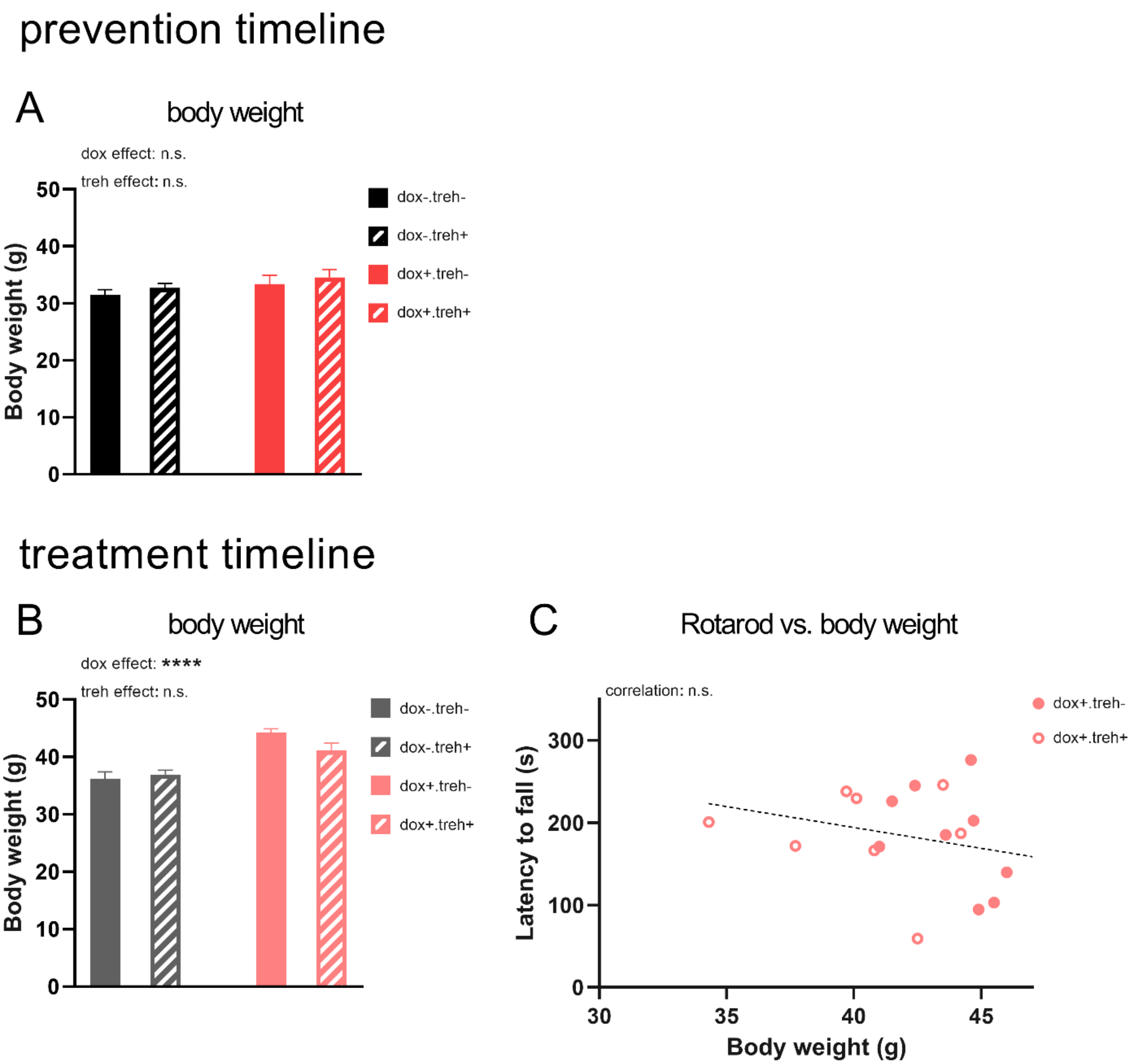


**Supplementary Figure 2. Changes in body weight did not alter the performance on the Rotarod**

**A.** Neither dox nor trehalose had any effect on the body weights of the mice under the prevention timeline. B. Under the treatment timeline, dox-induced mice were significantly heavier than their counterparts. No such effect was observed for trehalose supplemented mice. C. The increase in body weight in the dox-induced mice did not correlate with any changes in performance during the Rotarod acceleration test. Data presented as mean ±S.E.M. ****$p<0.0001$, $^{n.s.}p>0.05$.
